# Increased GITRL impairs the function of MDSCs and exacerbates primary Sjögren’s syndrome

**DOI:** 10.1101/373753

**Authors:** Jie Tian, Ke Rui, Yue Hong, Xiang Lin, Xiaohui Wang, Fan Xiao, Huaxi Xu, Liwei Lu, Shengjun Wang

**Author notes:** Jie Tian and Ke Rui contributed equally to this work. **Address correspondence to:** Dr. Shengjun Wang, Department of Laboratory Medicine, The Affiliated People’s Hospital, Jiangsu University, Zhenjiang, 212013, China.; Dr. Liwei Lu, Department of Pathology and Center of Infection and Immunology, The University of Hong Kong, Pokfulam Road, Hong Kong, China.

## Abstract

It is largely unclear how MDSCs contribute to the development of primary Sjögren’s syndrome (pSS). In experimental SS (ESS) mice, MDSCs were significantly increased but exhibited gradually diminished suppressive capacity during the disease progression. The ligand for glucocorticoid-induced TNFR family-related protein (GITRL) was increased with the development of pSS, and the increased GITRL was found to down-regulate the function of MDSCs while blocking GITR signal in MDSCs significantly restored their function and ameliorated ESS progression in mice. In pSS patients, expanded MDSCs expressed lower level of arginase were observed in patients with higher SSDAI. Moreover, the increased GITRL in serum was also found to closely correlate with the aberrant function of MDSCs. Together, our studies have demonstrated a critical role of GITRL in modulating the suppressive capacity of MDSCs in pSS, which may facilitate the validation of GITRL as a therapeutic target for the treatment of pSS.

## Introduction

Primary Sjögren’s syndrome (pSS), a systemic autoimmune disease, is characterized by lymphocytic infiltration of the exocrine glands, primarily salivary and lachrymal glands, leading to the loss of secretary function (Fox, 2005; Nocturne & Mariette, 2013). Although the etiology of pSS remains largely unclear, many studies have demonstrated the involvement of dendritic cells (DCs), NK cells, B cells and T cells in the pathogenesis of SS (Adamson *et al*, 1983; Cornec *et al*, 2012; Nocturne & Mariette, 2013; Singh & Cohen, 2012). Recent studies have found that Th1 and Th17 immune responses play a critical role in the development of pSS (Kolkowski *et al*, 1999; Lin *et al*, 2015; Mitsias *et al*, 2002; Nguyen *et al*, 2008; Sakai *et al*, 2008; Xiao *et al*, 2017).

Myeloid-derived suppressor cells (MDSCs) are a heterogeneous population of immature myeloid cells with immunosuppressive functions (Gabrilovich & Nagaraj, 2009). Murine MDSCs are characterized by co-expression of CD11b and Gr-1, which can be further subdivided into CD11b^+^Ly-6G^-^Ly-6C^high^ monocytic MDSCs (M-MDSCs) and CD11b^+^Ly-6G^+^Ly-6C^low^ polymorphonuclear MDSCs (PMN-MDSCs). Human MDSCs are CD11b^+^CD33^+^HLA-DR^−^ cells. MDSCs have been shown to promote tumor progression by suppressing T cell-mediated anti-tumor immunity in cancer (Nagaraj *et al*, 2009; Rabinovich *et al*, 2007). Recently, MDSCs are shown to be involved in the pathogenesis of various autoimmune disorders, including type 1 diabetes, multiple sclerosis (MS), rheumatoid arthritis (RA) and systemic lupus erythematosus (SLE), but the role of MDSCs in the development of pSS has remained poorly understood.

Glucocorticoid-induced tumor necrosis factor receptor (TNFR) family-related protein (GITR) is a type I transmembrane protein of TNF superfamily, which is highly expressed on Tregs, but low levels on conventional effector T cells and is rapidly up-regulated after activation (Kanamaru *et al*, 2004). The ligand for GITR (GITRL) is a type II transmembrane protein predominantly expressed on endothelial cells, dendritic cells (DCs), macrophages and B cells (Kim *et al*, 2003; Tone *et al*, 2003). The interaction of GITR and GITRL has been demonstrated to modulate both innate and adaptive immune responses (Azuma, 2010). The engagement of GITR on effector T cells exhibits a positive co-stimulatory signal leading to T cell proliferation and cytokine production (Chattopadhyay *et al*, 2008). However, GITR stimulation on regulatory T cells has been suggested to abolish the suppressive capacity of the cells (Chattopadhyay *et al*, 2007; Shevach & Stephens, 2006). Previous studies including our recent findings have revealed that GITR/GITRL pathway is involved in the pathogenesis of autoimmune diseases, including RA (Cuzzocrea *et al*, 2005; Wang *et al*, 2012), MS (Kohm *et al*, 2004) and autoimmune diabetes (You *et al*, 2009). Recently, GITRL is reported to be closely associated with the disease severity in pSS patients and MRL*-Fas^lpr^* mice (Gan *et al*, 2013; Saito *et al*, 2013). However, the underlying mechanism for GITRL in regulating the progression in pSS has been largely unclear.

In this study, we characterized the kinetic change and functional alteration of MDSCs in experimental SS (ESS) mice and pSS patients and identified a critical role of GITRL in regulating the suppressive function of MDSCs in the pathogenesis of pSS.

## Results

### Expanded MDSCs during ESS development in mice

In ESS mice, the kinetic change of CD11b^+^Gr-1^+^ MDSCs was analyzed in the spleen, CLN and peripheral blood during the progression of the disease. Compared to the naïve mice (day 0) or control mice immunized with adjuvant alone, percentages of MDSCs in different samples remarkably increased and reached a peak around day 14 or 35 post immunization (Figure 1A-C). Flow cytometric analysis identified a population of MDSCs infiltrated in SG 10 weeks after immunization (Figure 1D), which was further confirmed by immunofluorescence microscopy (Figure 1E). Similarly, both the number and the proportion of two subsets, CD11b^+^Ly-6G^-^Ly-6C^high^ M-MDSCs and CD11b^+^Ly-6G^+^Ly-6C^low^ PMN-MDSCs were also increased during the course of ESS (Figure 1F, G). Of note, mice treated with adjuvant showed only a slight increase of MDSCs at the early stage.

**Figure 1.**
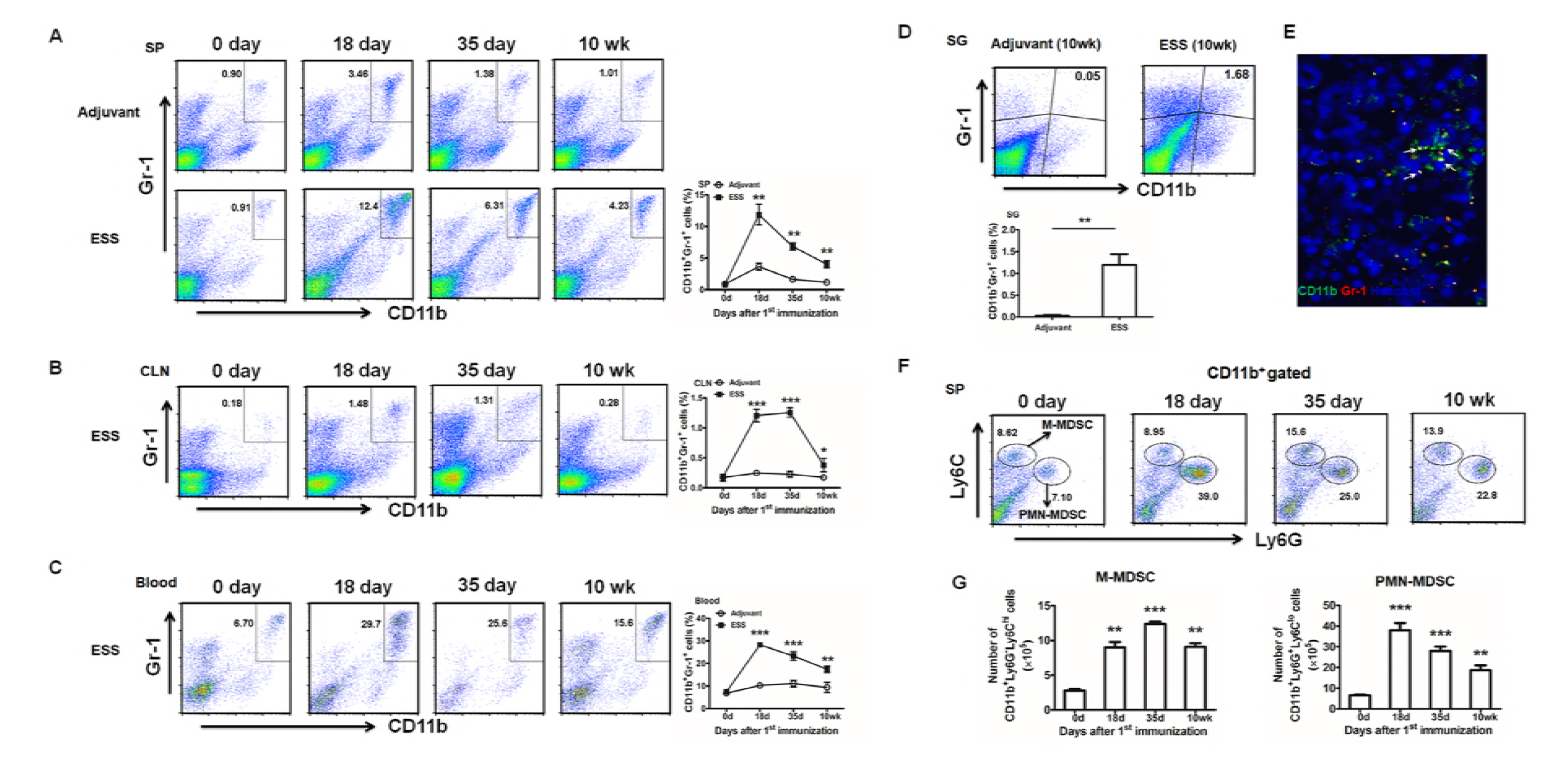
Expansion of MDSCs during the progression of ESS. (A-C) Kinetic changes of CD11b^+^Gr-1^+^ MDSCs in spleens (SP) (A), cervical lymph nodes (CLN) (B) and blood (C) during ESS development (n=12/group). (D, E) MDSCs in SG from ESS mice 10 weeks post immunization were analyzed by flow cytometry (D) and immunofluorescence microscopy (E),CD11b^+^Gr-1^+^ MDSCs (arrows) in the submandibular gland were stained with FITC-labeled anti-CD11b mAb and PE-labeled anti-Gr-1 mAb (original magnification×200) (n=6/group). (F, G) Both frequencies and absolute numbers of CD11b^+^Ly6G^-^Ly6C^hi^ (M-MDSCs) and CD11b^+^Ly6G^+^Ly6C^lo^ (PMN-MDSCs) in spleens were detected during ESS development (n=12/group). Data are shown as mean± SD of three independent experiments. ***p< 0.001, **p<0.01, *p<0.05.

### MDSCs gradually lose their suppressive capacity during ESS progression

We next defined the maturation and function of the increased MDSCs, the expression of CD40, CD80, CD86 and MHCII on MDSCs and their suppressive capacity on T cell proliferation were analyzed. As shown in Figure 2A, MDSCs (M-MDSCs/PMN-MDSCs) isolated from the early stage of ESS (day 18) expressed low levels of CD40, CD80, CD86 and MHCII, which displayed an immature and undifferentiated phenotype. However, expression of these molecules was gradually up-regulated with the development of the disease, and MDSCs from the late stage of ESS (10 weeks) expressed high levels of these surface markers, which exhibited a more mature status. Co-culture of MDSCs (M-MDSCs/PMN-MDSCs) and T cells indicated the early-stage MDSCs displayed highly efficient activity in suppressing the T cell proliferation while the late-stage cells almost lost their suppressive capacity (Figure 2B). Similarly, the suppressive factors secreted by MDSCs, including arginase and NO, were also down-regulated with the development of ESS (Figure 2C). Furthermore, MDSCs infiltrated in SG produced less arginase when compared to the early-stage MDSCs in spleens, which indicates that MDSCs migrated into the local inflammatory sites are of weak suppressivity (Figure 2D). Together, these results suggest that the suppressive capacity of MDSCs was down-regulated during the progression of ESS.

**Figure 2.**
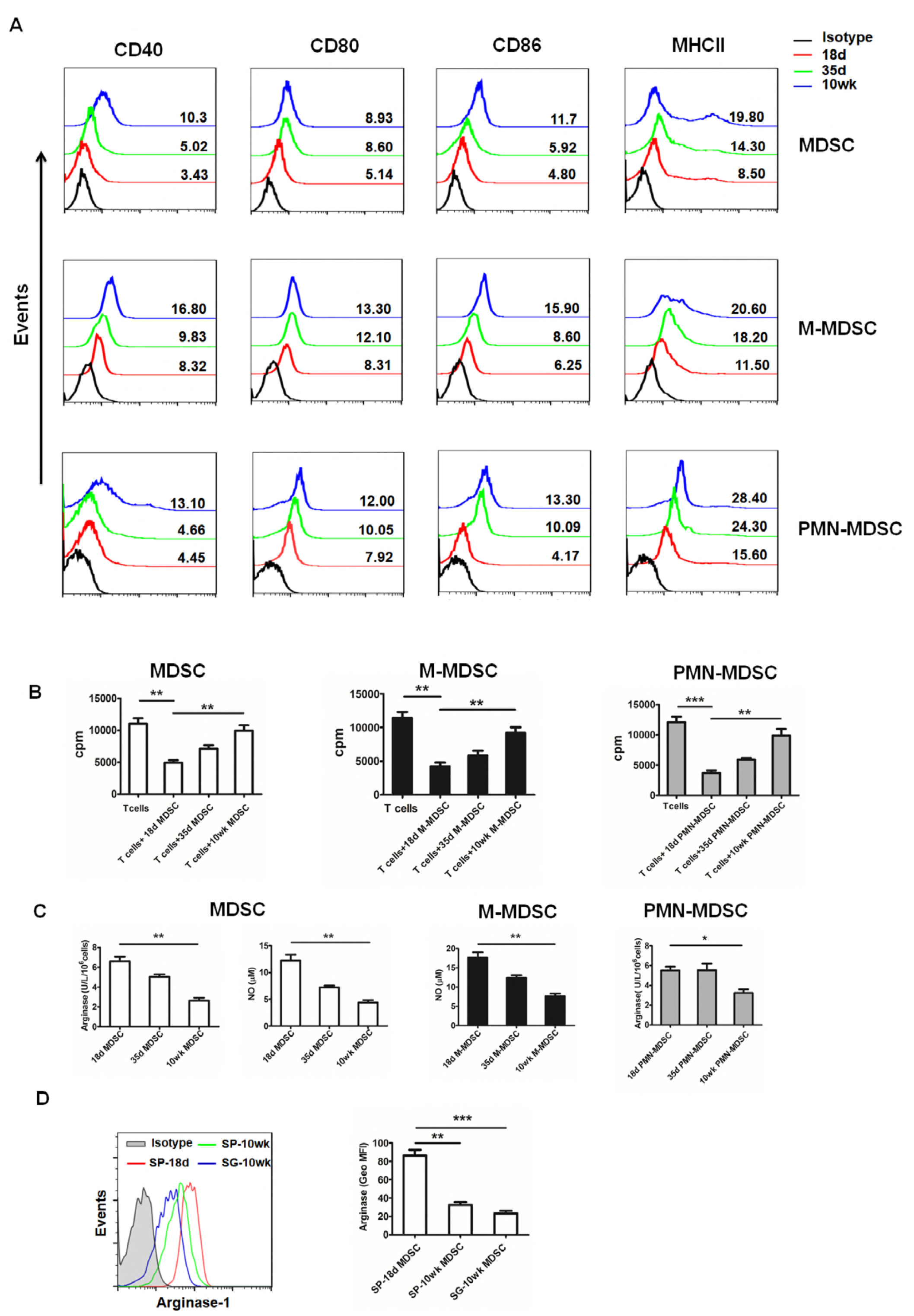
Impaired suppressive capacity of MDSCs during the development of ESS. Total MDSCs, M-MDSCs and PMN-MDSCs were isolated from spleens of ESS mice at different time points (18 days, 35 days and 10 weeks after first immunization). (A) Cells were stained with specific Abs against CD40, CD80, CD86 and MHCII. (B) Total MDSCs, M-MDSCs and PMN-MDSCs at different time points were co-cultured with CD4^+^ T cells in the presence of anti-CD3 mAb and anti-CD28 mAb for 72 h. Suppressivity on T cell proliferation was measured by incorporation of [^3^H]-thymidine. (C) Arginase activity and NO levels released by MDSCs were measured as described in Methods. (D) Cells in spleens (18 days/10 weeks) and submandibular glands (10 weeks) were subjected to analyze the arginase expression of MDSCs by flow cytometry. Data are shown as mean± SD of three independent experiments. ***p< 0.001, **p<0.01, *p<0.05.

Next, we further observed the suppressivity of MDSCs in vivo. Early-stage or late-stage MDSCs were adoptively transferred to ESS mice on days 18 and 25 (Figure EV1A). Remarkably, the early-stage MDSCs, which possessed strong suppressive capacity, significantly reversed the saliva flow rate to normal level while the late-stage cells did not (Figure EV1B). Furthermore, levels of autoantibodies against total SG antigens, ANA and anti-M3R antibodies were also decreased after early-stage MDSCs treatment (Figure EV1C). Notably, the early-stage MDSCs treated group displayed a smaller CLN and SG when compared to the control group (Figure EV1D). Histological examination further identified the pathological changes in local submandibular glands. Only little lymphocyte infiltration was observed in SG after early-aged MDSCs treatment. In contrast, the late-aged MDSCs treatment did not display any therapeutic effect, and even remarkably aggravated the tissue destruction, showing multiple lymphocytic foci in SG (Figure EV1E). Additionally, Th1 and Th17 responses were also reduced after the treatment of early-stage MDSCs while the late-stage MDSCs did not (Figure EV1 F, G). Similar results were also observed in the serum levels of IFN-γ and IL-17 (Figure EV1H). Taken together, these findings reveal that early-stage MDSCs could ameliorate the progression of ESS and reduce the Th1/Th17 responses while late-stage MDSCs could not.

### GITRL impairs the function of MDSCs and enhances the progression of ESS

The above in vitro and in vivo experiments have clarified the reduction of the suppressive function of MDSCs during the development of ESS, we next investigated the potential factors responsible for the altered function of MDSCs. We firstly found that GITR was expressed on total MDSCs and two subpopulations (Figure 3A). Moreover, the level of GITR was up-regulated with the disease progression (Figure 3B). Notably, GITRL was also gradually increased in the spleen, CLN and SG with the progression of the disease (Figure 3C). To further ascertain the contributed cell population, we analyzed three populations which mainly expressed GITRL in peripheral immune organs. As shown in Figure 3D and 3E, GITRL expressed on both macrophages and B cells was increased in ESS mice. Thus, these results suggest that MDSCs in ESS mice might be regulated by the increased GITRL in vivo.

**Figure 3.**
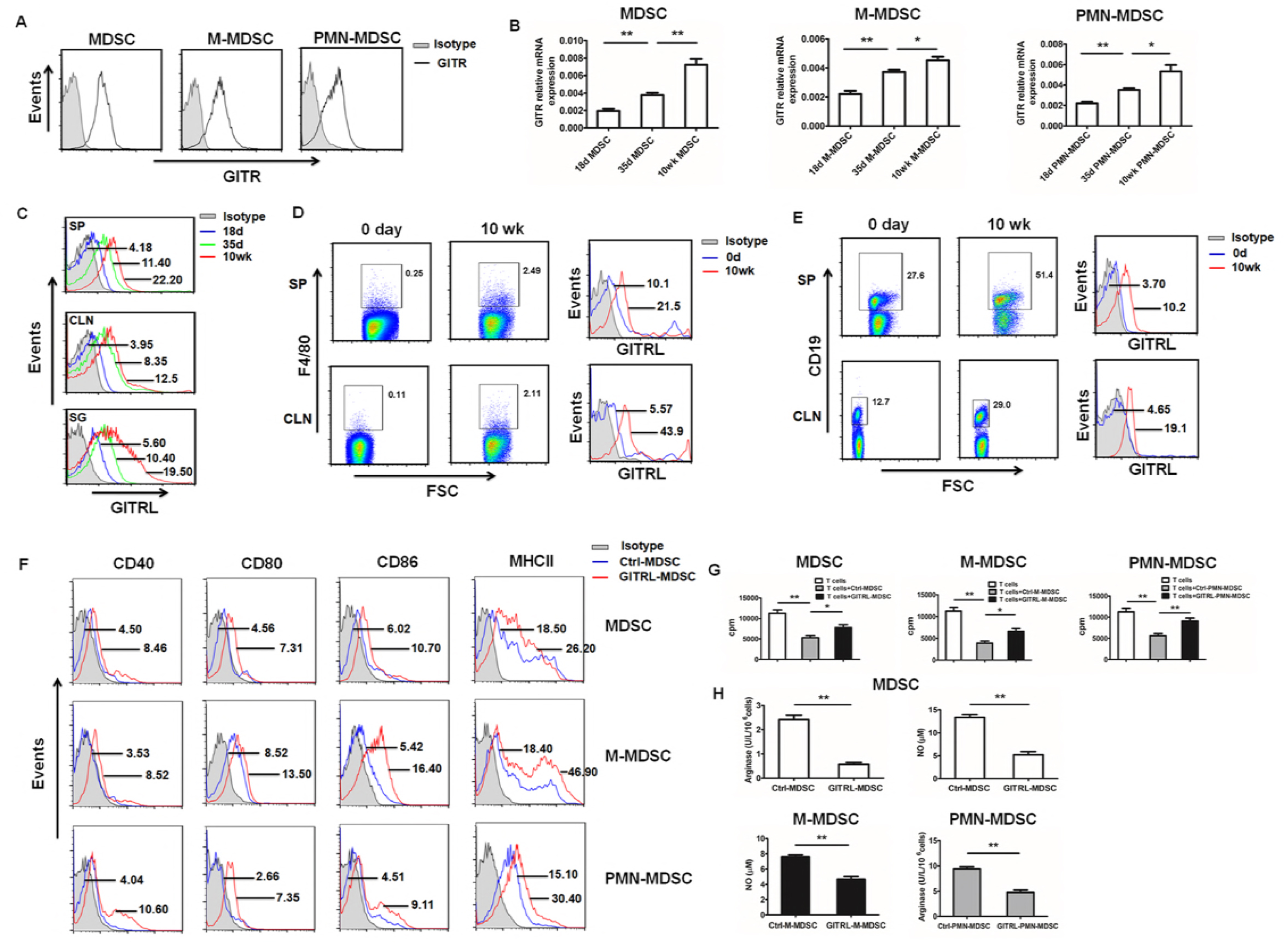
Increased GITRL promotes the maturation and down-regulates the suppressive capacity of MDSCs. (A) Splenic total MDSCs, M-MDSCs and PMN-MDSCs were isolated from ESS mice on day 18 after the first immunization, and FCM was used to analyze the GITR expression (n=12). (B) qRT-PCR was used to analyze the mRNA expression of GITR in MDSCs and their subsets on day 18, day 35 and 10 weeks after first immunization (n=12). (C) Protein levels of GITRL were analyzed in spleen, CLN, SG isolated on day 18, day 35 and 10 weeks after first immunization (n=12). (D, E) Frequencies of F4/80^+^ macrophages (D) and CD19^+^ B cells (E) in spleens and cervical lymph nodes on day 0 and 10 weeks after first immunization were analyzed using flow cytometry, and GITRL expression on macrophages and B cells were further detected. (F-H) Isolated splenic MDSCs, M-MDSCs and PMN-MDSCs from ESS mice on day 18 after the first immunization were treated with GITRL (5μg/mL) or control protein for 48h, and 0.2ng/mL GM-CSF was added to sustain the survival of MDSCs. (F) Levels of CD40, CD80, CD86 and MHCII on MDSCs (M-MDSCs/PMN-MDSCs) treated with GITRL or control protein were measured by flow cytometry. (G) GITRL-treated MDSCs were collected to co-culture with CD4^+^T cells (MDSC: T cell ratio of 1:1) in the presence of anti-CD3 mAb and anti-CD28 mAb for 72 h. Suppression of T-cell proliferation was measured by incorporation of [^3^H]-thymidine. (H) Arginase activity and NO levels released by MDSCs and their subsets were measured. Data are shown as mean± SD of three independent experiments. **p < 0.01, *p < 0.05.

To investigate whether the function of MDSCs would be regulated by the increased GITRL, recombinant GITRL was used to stimulate the early-stage MDSCs isolated from ESS mice. After GITRL treatment, levels of CD40, CD80, CD86 and MHCII on total MDSCs, M-MDSCs and PMN-MDSCs were enhanced (Figure 3F), and the suppressive function of MDSCs (M-MDSCs and PMN-MDSCs) on CD4^+^ T cells was decreased (Figure 3G). Similarly, reduced arginase and NO was observed after GITRL stimulation (Figure 3H). All these results suggest GITRL could promote the maturation of MDSCs and down-regulate the suppressive capability of the cells.

We next investigated the effect of GITRL-treated MDSCs in inhibiting the ESS development. Early-stage MDSCs treated with or without GITRL were adoptively transferred to ESS mice on days 18 and 25 respectively (Figure EV2A). When compared to the Ctrl-MDSC group, saliva flow rate was significantly decreased and the levels of autoantibodies were enhanced after transferring with GITRL-treated MDSCs (Figure EV2B-E). In addition, GITRL-MDSCs treatment showed a bigger CLN and SG, and more lymphocytes were detected in local submandibular glands (Figure EV2F, G). Furthermore, GITRL-MDSCs treatment did not reduce the Th1 and Th17 responses while the Ctrl-MDSCs significantly inhibit the immune responses (Figure EV2H-J). Together, these findings suggest that GITRL treatment make the early-stage MDSCs lose their primary capability in inhibiting the ESS development.

### GITRL treatment attenuates the suppressive effect of MDSCs in ESS mice

Exogenous GITRL was administrated into the ESS mice starting from day 18 for three times (Figure 4 A), and the GITRL treatment significantly exacerbated the disease development. The reduced saliva flow rate, enhanced autoantibodies as well as more serious lymphocyte infiltration in submandibular glands were observed (Figure 4 B-E). Then, the maturation and the function of MDSCs in vivo was observed. As shown in Figure 4F, the expression of co-stimulatory molecules and MHCII on MDSCs (M-MDSCs/PMN-MDSCs) was enhanced with varying degrees, and the suppressive capacity of MDSCs in ESS mice was inhibited after GITRL administration (Figure 4G, H).

**Figure 4.**
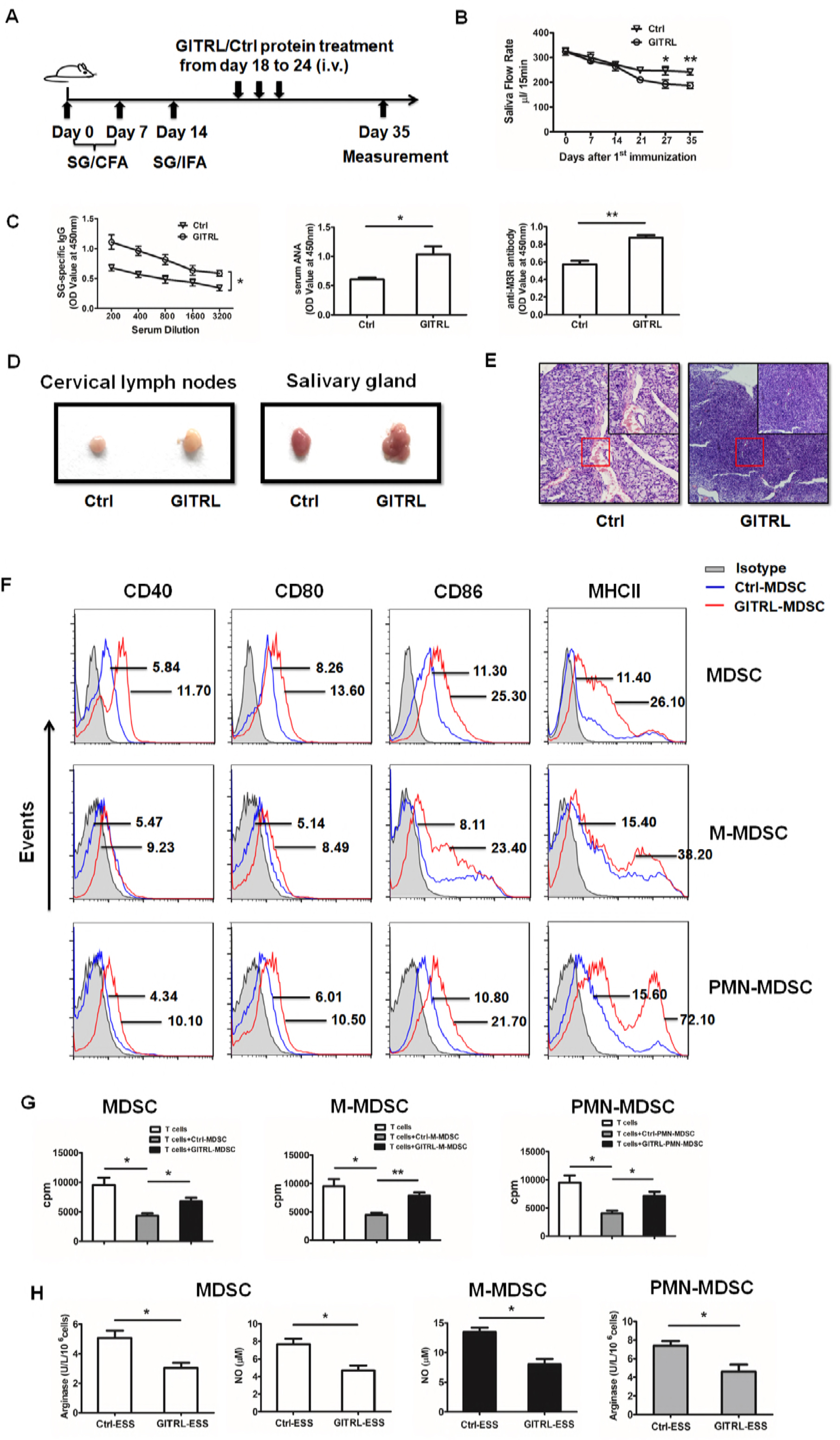
GITRL modifies the suppressive capacity of MDSCs in ESS mice. (A) Graphic scheme of ESS induction and GITRL administration. Recombinant GITRL or control protein (2mg/kg) was intravenously administered on days 18, 21 and 24. Mice were sacrificed on day 35 (n=12/group). (B) The saliva flow rates were measured in GITRL treated group and the control group. (C) Autoantibodies against SG antigens, ANA and anti-M3R antibodies were analyzed in the serum of mice with different treatment. (D) Representative micrographs show the sizes of CLN and SG from two groups. (E) ESS mice were injected with GITRL three times one week for 3 weeks, starting at 18 days after the first immunization. The histological evaluation of glandular destruction in each group was performed on tissue sections of submandibular glands with H&E staining 15 weeks post first immunization. (F) Splenic MDSCs, M-MDSCs and PMN-MDSCs from two groups were isolated and the expression of CD40, CD80, CD86 and MHCII on MDSCs was measured by flow cytometry. (G) Isolated MDSCs (M-MDSCs/PMN-MDSCs) were collected to co-culture with CD4^+^T cells (MDSC: T cell ratio of 1:1) in the presence of anti-CD3 mAb and anti-CD28 mAb for 72 h. Suppression of T-cell proliferation was measured by incorporation of [^3^H]-thymidine. (H) Arginase activity and NO levels released by MDSCs and their subsets were measured. Data are shown as mean± SD of three independent experiments. **p < 0.01, *p < 0.05.

### Blocking GITR signals in MDSCs restores their capacity in suppressing ESS development

To further clarify whether the altered function of MDSCs in vivo was regulated by the engagement of GITR, we adoptively transferred the MDSCs interfered with GITR into ESS mice on day 32 (Figure 5A). The previous data have shown that the level of GITRL in vivo have remarkably increased around 35 days after immunization. Thus, we assumed the function of transferred MDSCs would be regulated under the environment with high concentration of GITRL. Consistent with our hypothesis, the therapeutic effect of MDSCs in inhibiting ESS was extremely worse. However, MDSCs interfered with GITR showed efficient therapeutic effect. The saliva flow rate was recovered to normal level and the secretion of autoantibodies was significantly reduced in siGITR-MDSCs group (Figure 5B-E). Also, when compared to the Ctrl-MDSCs and ESS group, the size of CLN and SG was smaller, and fewer lymphocytes were infiltrated in submandibular glands in siGITR-MDSCs group (Figure 5F, G). Furthermore, flow cytometric analysis detected reduced Th1 and Th17 responses in ESS mice treated with siGITR-MDSCs (Figure 5H-J). All these data imply that the suppressive capacity of MDSCs in controlling the progress of ESS was regulated by GITR/GITRL pathway.

**Figure 5.**
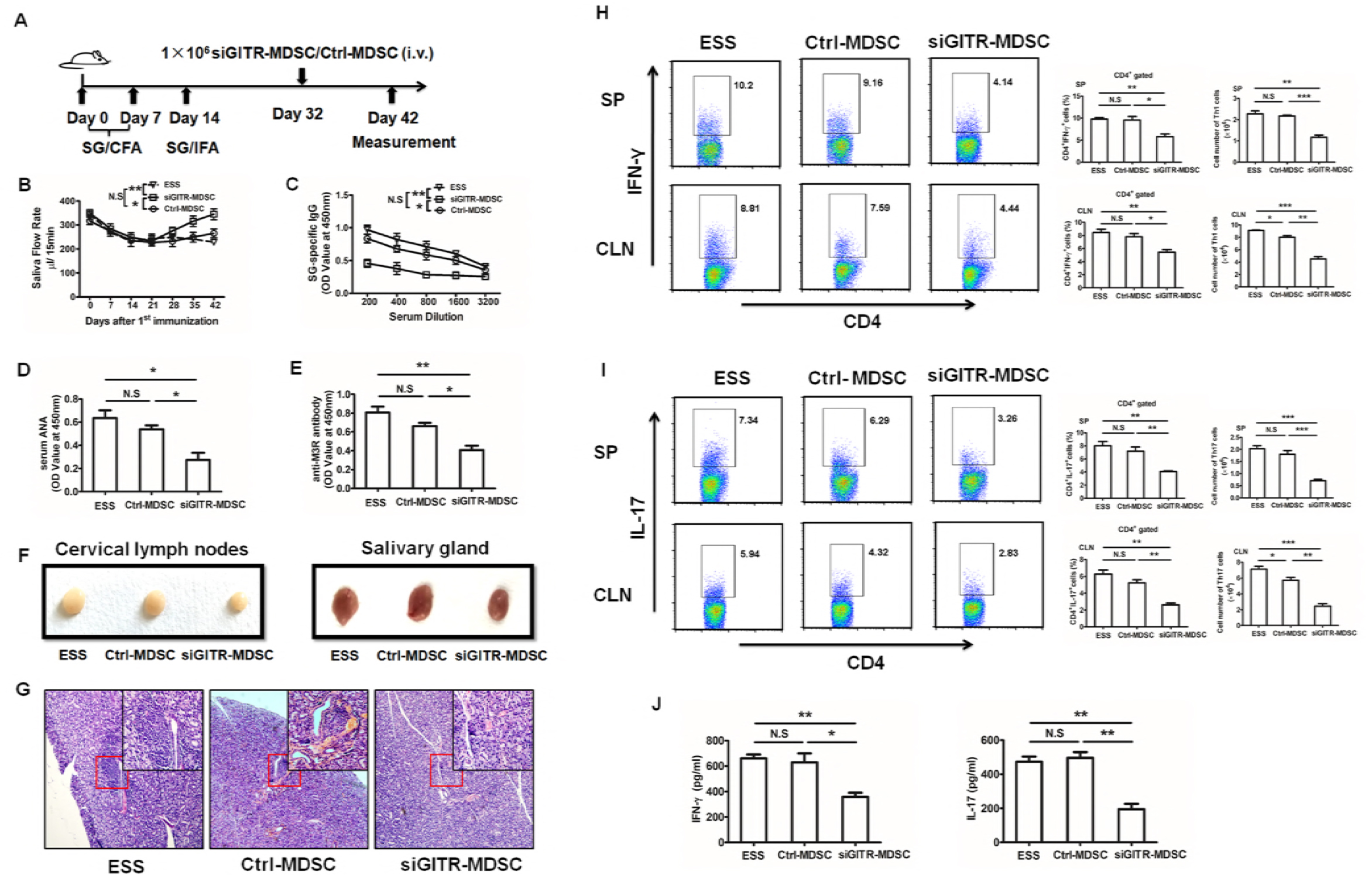
MDSCs interfered with GITR recover their capacity in inhibiting ESS development. (A) Graphic scheme of ESS induction and MDSC administration. MDSCs isolated from spleens of ESS mice on day 18 were transfected with GITR siRNA (siGITR) or negative control for 24 h, and then 1×10^6^ siGITR-MDSCs or Ctrl-MDSCs were intravenously injected on day 32 after the first immunization. Mice were sacrificed on day 42 (n=12/group). (B) The saliva flow rates were measured in each group. (C-E) Autoantibodies against SG antigens (C), ANA (D) and anti-M3R antibodies (E) were analyzed in the serum of mice with different treatment. (F) Representative micrographs show the sizes of CLN and SG from different groups. (G) ESS mice were transferred with different MDSCs once for 3 weeks, starting at 32 days post the first immunization. The histological evaluation of glandular destruction in each group was performed on tissue sections of submandibular glands with H&E staining 15 weeks post first immunization. (H, I) Both proportions and numbers of CD4^+^IFN-γ^+^ Th1 cells (H) and CD4^+^IL-17^+^ Th17 cells (I) were measured in SP and CLN of mice with different treatment on day 42. (J) Serum levels of IFN-γ and IL-17 were detected in different groups on day 42. Data are shown as mean± SD of three independent experiments. ***p < 0.001, **p < 0.01, *p < 0.05. N.S, no significance.

### Increased GITRL correlates with impaired function of MDSCs in pSS patients

Flow cytometric analysis was performed to examine the frequencies of CD11b^+^CD33^+^HLA-DR^-^ MDSCs in peripheral blood from pSS patients and healthy controls. Compared to healthy controls, the percentage of MDSCs was significantly increased in patients with pSS (Figure 6A). A further analysis showed these accumulated MDSCs in pSS patients secreted low level of arginase (Figure 6B), and the level of arginase was negatively correlated with the disease activity (Figure 6C). In pSS patients, both the serum level of GITRL and GITR expression on MDSCs were significantly enhanced and displayed positive correlations with the activity of the disease (Figure 6D-G). Moreover, arginase in MDSCs from pSS patients was negatively correlated with GITRL and GITR (Figure 6H-I). To further explore the regulation of GITRL on the function of human MDSCs, MDSCs induced from healthy donors were treated with recombinant hGITRL. Similar to the data in mice, the suppressive capacity of MDSCs on CD4^+^ T cells was significantly decreased after GITRL treatment (Figure 6J), and the arginase level in MDSCs was also reduced (Figure 6K). All these data suggest that the suppressive function of MDSCs in pSS patients is closely related to the severity of the disease, which is probably regulated by GITR/GITRL pathway.

**Figure 6.**
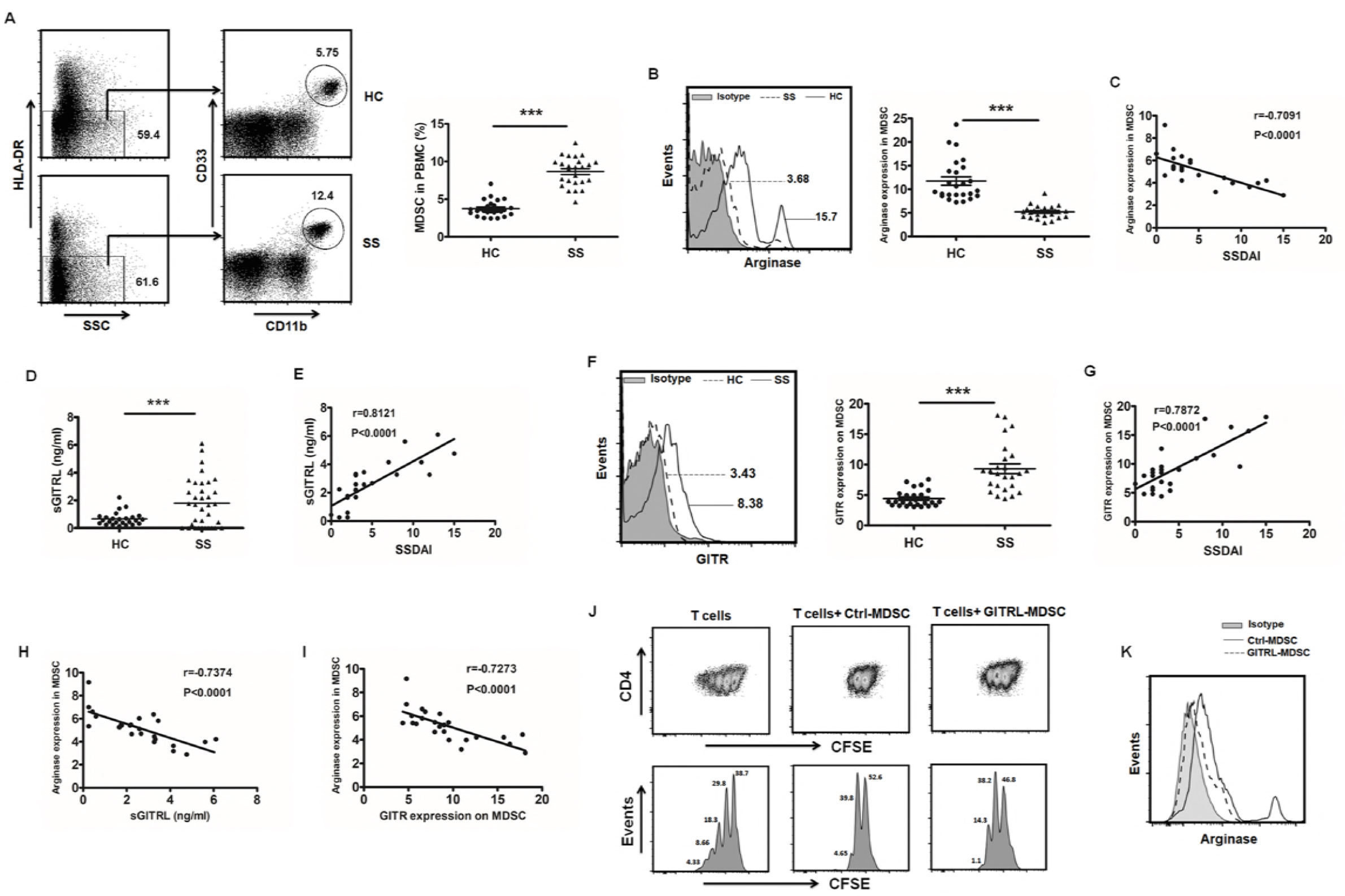
Increased GITRL displays negative correlation with the suppressive function of MDSCs in pSS patients. (A) CD11b^+^CD33^+^HLA-DR^-^ MDSCs in blood was measured in pSS or healthy controls (n=25), the representative flow cytometry graph and statistical graph were shown respectively. (B, C) The level of arginase in MDSCs from two groups was detected by flow cytometry (B), and its relationship with SSDAI was analyzed (C). (D, E) Plasma from pSS patients and healthy controls were collected to detect GITRL levels (D), and a correlation between GITRL and SSDAI was analyzed (E). (F) GITR expression on MDSCs from pSS or healthy controls was observed, and the representative flow chart and the statistical results were shown. (G) The correlation between GITR and SSDAI was observed. (H, I) Correlations between arginase and GITRL (H), GITR (I) were identified. (J, K) MDSCs were induced by PBMCs supplemented with GM-CSF and IL-6, and then CD33^+^ cells were isolated to be stimulated with hGITRL or control protein for 48h. Cells were harvested to coculture with CFSE-labeled autologous CD4^+^T cells under CD3/CD28 stimulation. After 3 days, the T cell proliferation was determined by flow cytometric analysis of CFSE dilution (J). Arginase expression in MDSCs was detected by flow cytometry (K). Results are expressed as mean± SD. The data are representative of three independent experiments. ***p < 0.001.

### Discussion

Due to the suppressive capability of MDSCs, accumulating investigations have devoted to exploit their potential in the treatment of autoimmune diseases (Fujii *et al*, 2013; Ioannou *et al*, 2012; Jeong *et al*, 2018; Li *et al*, 2014; Zhu *et al*, 2007). However, in this study, we showed that frequencies of MDSCs and their subsets were enhanced, but they gradually lost their suppressive function with the development of ESS, thus leading to the uncontrollable expansion of Th1 and Th17 immune responses. Further investigation demonstrated that it was the enhanced GITRL that modulate the suppressivity of MDSCs directly through GITR/GITRL pathway. Taken together, our findings have revealed that the increased GITRL could promote the maturation of MDSCs, and down-regulated the suppressive capacity of the cells, which further promoted the progression of pSS.

To date, there was no evidence showing the relationship between MDSCs and pSS. However, the role of MDSCs in other autoimmune diseases has been discussed for years. The expansion of MDSCs was found in the majority of the autoimmune diseases, such as MS, RA, type I diabetes, autoimmune uveoretinitis and SLE (Fujii *et al*, 2013; Guo *et al*, 2016; Ioannou *et al*, 2012; Jeong *et al*, 2018; Yi *et al*, 2012; Yin *et al*, 2010; Zhu *et al*, 2007). Nevertheless, the conclusions about the role of MDSCs in different autoimmune diseases were not totally identical, and even in the same disease, the effect of MDSCs could be opposite. In the EAE mouse model, Ioannou *et al.* found that only G-MDSCs subpopulation was increased and adoptive transfer of G-MDSCs isolated from EAE mice significantly suppressed the development of EAE and inhibited Th1 and Th17 immune responses (Ioannou *et al*, 2012). However, Yi *et al.* demonstrated that M-MDSCs in EAE could promote the differentiation of Th17, and depletion of MDSCs remarkably reduced the Th17 response and ameliorated the severity of EAE (Yi *et al*, 2012). The similar conflicting results were also observed in the autoimmune arthritis. Fujii *et al.* suggested a protective role of G-MDSCs in collagen-induced arthritis (CIA) mice (Fujii *et al*, 2013) while Guo *et al.* reported M-MDSC in CIA mice could promote Th17 cell polarization and then promote the disease progression (Boros *et al*, 2016). The contrary conclusions might be affected by multiple factors, such as different pathogenesis of diseases, various inner environments, different MDSC subsets and mouse genetic backgrounds. Owing to the extraordinary heterogeneity and plasticity in phenotypes and functions, myeloid cell infiltrated at different stages of a disease contributes to a differential role in the disease promotion and amelioration. We first found the increased MDSCs in pSS patients with higher SSDAI expressed lower level of arginase. In the ESS mouse model, MDSCs were also significantly enhanced in different tissues. However, the function and the phenotype of MDSCs infiltrated in different stages were altered. The early-stage MDSCs displayed strong suppressive capacity while the late-stage cells almost lost their inhibition on T cells. Furthermore, levels of CD40, CD80, CD86 and MHCII were increased on the late-stage MDSCs, which exhibited a more mature phenotype of myeloid cells. Intriguingly, CD11b^+^Gr-1^+^ cells infiltrated in SG expressed low level of arginase, suggesting that MDSCs in local tissues are not associated with immunosuppression, but probably induce or promote the local inflammation, and further investigation is needed to figure out this issue. Additionally, adoptive transfer of early-stage MDSCs to ESS mice significantly ameliorated the severity of ESS, showing an obvious therapeutic effect on the disease. However, the late-stage MDSCs showed no ability to alleviate the disease. Interestingly, more serious lymphocyte infiltration and the higher Th1, Th17 immune responses suggest that the late-stage MDSCs might have the ability to promote the inflammation, and we further found that the late-stage MDSCs could induce the differentiation of Th17 cells in vitro (Unpublished data). All these data suggest that the microenvironment of different stages would modify the status of MDSCs, which will determine their property of proinflammation or suppression. Therefore, in order to explore effective MDSC-based therapies, we must first clarify the phenotype, function and differentiation of MDSCs in the disease, and figure out the key factors in different inflammatory environments that affect the potency of MDSCs.

The conflicting results on the role of MDSCs in preventing autoimmune diseases are mainly due to the high heterogeneity and plasticity of myeloid cells whose phenotypes and functions are largely dependent on local microenvironments (Cripps & Gorham, 2011; Gabrilovich & Nagaraj, 2009; Melero-Jerez *et al*, 2016; Sunderkotter *et al*, 2004; Youn & Gabrilovich, 2010). In this study, we found that GITR was expressed on MDSCs for the first time, both in ESS mice and pSS patients, and the expression of GITR on MDSCs was significantly enhanced with the progression of the disease. Besides, GITRL was also significantly increased in ESS mice and patients. Actually, Gan *et al.* have reported the increased GITRL was closely associated with the severity of the disease in pSS patients (Gan *et al*, 2013), which was consistent with our results. Further investigation revealed that the increased GITRL was mainly ascribed to macrophages and B cells. Thus, the next question is whether MDSCs with suppressive function will be regulated by the increased GITRL. Then, we utilized the recombinant GITRL to stimulate MDSCs in vitro and administered it into ESS mice in vivo, the result showed GITRL could down-regulate the suppressive effect of MDSCs and promote them differentiate into a mature phenotype in vitro and in vivo. The similar results were also observed in humans, GITRL could impair the suppressivity of induced MDSCs in humans. Moreover, the early-stage MDSCs modified by GITRL lost their primary capacity to ameliorate the ESS development. In the environment with high level of GITRL, MDSCs interfered with GITR exhibited efficient therapeutic effect in vivo, indicating the GITR/GITRL pathway is the predominant element for the regulation of MDSCs. Additionally, consistent with the late-stage MDSCs, adoptive transfer of GITRL-treated MDSCs could also lead to slight enhancement of Th1 and Th17 cell responses when compared to the ESS mice, suggesting their potential proinflammatory effect, and further studies are needed to clarify this issue. Together, all these data suggest, as cells with high heterogeneity and plasticity, MDSCs can not only lose their suppressive capacity after encountering inflammatory environments, but also can convert into pathogenic cells that might accelerate the inflammatory process in autoimmune diseases.

In conclusion, our findings demonstrated the increased GITRL modulated the phenotype and the function of MDSCs, thus inducing the progression of pSS. Our data noted that MDSCs were not terminally differentiated cells, the microenvironment can modulate them and direct their proinflammatory or suppressive phenotypes. Therefore, to develop effective MDSC-based therapies, the plasticity of immunoregulation by MDSCs should be considered. Learning how to control the microenvironment may provide an important novel approach for better therapy in pSS.

## Materials and Methods

### Mice

Female C57BL/6 mice at 8-week-old were purchased from Experimental Animal Center of Yangzhou University. Mice were housed in a specific pathogen-free animal facility and all the experiments were approved by the Committee on the Use of Live Animals in Research and Teaching of Jiangsu University.

### Induction of ESS model

The ESS mouse model was induced as previously described (Lin *et al*, 2015). Briefly, bilateral salivary glands were isolated from female C57BL/6 mice for homogenization in PBS to prepare salivary glands (SG) proteins. Naïve mice were immunized with SG proteins emulsified in an equal volume of CFA (Sigma-Aldrich) to a concentration of 2 mg/mL (100μl each mouse) s.c. on the neck on days 0 and 7. On day 14, the booster injection was performed with a dose of 1 mg/mL SG proteins emulsified in Freund’s incomplete adjuvant (Sigma-Aldrich). Naïve mice immunized with adjuvant alone served as adjuvant controls.

### Detection of saliva flow rate

Saliva flow rates were measured as previously described (Lin *et al*, 2015). In brief, mice were anesthetized and injected intraperitoneally with pilocarpine (Sigma-Aldrich) at a dose of 5 mg/kg body weight. Saliva was then collected using a 20-μl pipet tip from the oral cavity for 15min.

### Flow cytometric analysis

For surface markers, single-cell suspensions were stained with relevant fluorochrome-conjugated mAbs: anti-mouse CD40, CD80, CD86, MHCII, GITR, GITRL, F4/80 and CD19 from eBioscience, anti-mouse CD11b, Gr-1, Ly6G and Ly6C from Biolegend, anti-mouse arginase-1 (R&D system); anti-human CD11b, HLA-DR, CD33, GITR, arginase-1 from Biolegend. For intracellular staining, cells were stimulated with PMA (Sigma-Aldrich, 50 ng/mL), ionomycin (Enzo, 1 μg/mL), monensin (Enzo, 2 μg/mL). After 5 h, cells were stained with anti-CD4 mAb (eBioscience), fixed, permeabilized, and stained with anti-IFN-γ mAb or anti-IL-17 mAb (eBioscience) according to the Intracellular Staining Kit (Invitrogen) instructions. Flow cytometry was performed using FACSCalibur Flow Cytometer (Becton Dickinson).

### MDSC isolation and suppression assay

CD11b^+^Gr-1^+^ MDSCs were isolated from the spleens of ESS mice using a FACSAria II SORP (Becton Dickinson) cell sorter. M-MDSCs and PMN-MDSCs were isolated using a mouse MDSC isolation kit (Miltenyi Biotec) following the manufacturer’s protocol. MDSC suppression assay was performed as we previously described (Tian *et al*, 2015).

### Quantitative real-time PCR (qRT-PCR)

The quantitative real-time PCR were performed as previously described (Tian *et al*, 2011). The sequences for the primers used are: mGITR, Forward −5’- CTCAGGAGAAGCACTATGGG’ −3’, Reverse-5’- AGCTGGGCAAGTCTTGTAG’ −3’. β-actin, Forward −5’-TGGAATCCTGTGGCATCCATGAAAC-3’, Reverse-5’-TAAAACGCAGCTCAGTAACAGTCCG-3’. β-actin was used as an internal control.

### Enzyme-linked immunosorbent assay

Autoantibodies against SG proteins, ANA (Elabscience), anti-M3 muscarinic receptor (M3R) antibodies, IFN-γ and IL-17 (eBioscience) in the serum of ESS mice and human GITRL in serum (R&D system) was assessed by ELISA.

### Detection of arginase activity and NO production

The activity of arginase and NO concentration were measured as previously described (Tian *et al*, 2013).

### Transfection

GITR siRNA, and negative controls were synthesized by RiboBio. Oligonucleotide transfection was performed with Entranster-R (Engreen Biosystem) according to the manufacturer’s instructions.

### Histological assessment and immunofluorescence microscopy

SG tissues were paraffin embedded, sectioned and stained with H&E. For immunofluorescence microscopy, frozen sections were incubated with FITC-conjugated anti-CD11b mAb (eBioscience) and PE-conjugated anti-CD33 mAb (eBioscience). Hoechst 33342 (Beyotime) was used to label nuclei. Matched isotype controls (eBioscience) were used for control staining.

### Patient samples

Whole blood samples were collected from patients with pSS (n=25) and healthy controls (n=25). All patients were diagnosed with active according to the SS Disease Activity Index (SSDAI) scores. The study protocols and consent forms were approved by the Institutional Medical Ethics Review Board of Jiangsu University.

### Generation of human MDSCs

Peripheral blood mononuclear cells (PBMC) were isolated from healthy volunteer donors by density gradient centrifugation using Ficoll-Hypaque solution (TBD sciences). PBMC were cultured with GM-CSF (10ng/ml, PeproTech) and IL-6 (10ng/ml, PeproTech) for 7 days, and the cytokines were refreshed every 2-3 days. After 1 week, all cells were collected and adherent cells were removed using nonprotease cell detachment solution Detachin (Genlantis). CD33^+^ cells were isolated using anti-CD33 magnetic microbeads (Miltenyi Biotec). The purity of isolated cells was >90%.

### T-cell suppression assay

Human CD4^+^ T cells were isolated from healthy volunteer donors using anti-CD4 microbeads (Miltenyi Biotec). CD4^+^ T cells were labeled with CFSE (5μM, Invitrogen) and co-cultured with human CD33^+^ cells at a ratio of 2:1 in 96-well plates (Costar) in the presence of anti-CD3 (eBioscience) and anti-CD28 (eBioscience) mAbs for 3 days. CFSE fluorescence intensity was analyzed to determine the proliferation of CD4^+^ T cells by flow cytometry.

### Statistical analysis

The statistical significance was determined by the Student’s t test or one-way ANOVA. Correlations were determined by a Spearman correlation coefficient. All analyses were performed using SPSS 16.0 software. p Values less than 0.05 were considered statistically significant.

## Acknowledgments

This work was supported by the Natural Science Foundation of Jiangsu (Grant Nos. BK20150533, BK20170563), National Natural Science Foundation of China (Grant Nos. 81601424, 81701612, 2014CB541904), Project funded by China Postdoctoral Science Foundation (Grant Nos. 2016M590423, 2017T100336), Summit of the Six Top Talents Program of Jiangsu Province (Grant No. 2017-YY-006), Young Talent Cultivation Program of Jiangsu University, Primary Research and Development Plan of Zhenjiang (Grant No. SH2017008).

## Author contributions

SW, LL, JT, KR and XL designed experiments, JT, KR, YH and XL performed experiments; JT, KR, YH, XL, XW and FX analysed the data. SW, LL and HX supervised the research. JT and KR wrote the manuscript.

